# The effects of elevated seawater pH and total alkalinity following dosing of sodium hydroxide in *Calanus finmarchicus*

**DOI:** 10.64898/2026.02.03.700700

**Authors:** Christopher S. Murray, Lukas Marx, Neelakanteswar Aluru, Zhaohui Aleck Wang, Ke Chen, Heather H. Kim, Anna Michel, Daniel C. McCorkle, Jennie E. Rheuban, Adam Suhbas

## Abstract

Ocean Alkalinity Enhancement (OAE) is a marine carbon dioxide removal (mCDR) strategy that involves adding alkaline substances to surface waters to enhance CO_2_ uptake and storage. The dispersal of alkaline materials such as sodium hydroxide (NaOH) into seawater can cause rapid increases in pH and total alkalinity (TA) that substantially exceeds natural variability in marine environments. Such fluctuations may negatively affect marine life, especially small animals like copepods who cannot avoid OAE plumes and whose physiological processes could be disrupted by large and rapid shifts in seawater pH. To address knowledge gaps regarding potential biological impacts of OAE, we studied these effects in *Calanus finmarchicus*, a keystone copepod species in the Northwest Atlantic Ocean. We exposed *C. finmarchicus* from the late juvenile copepodite stages and adult females to NaOH-dosed seawater at pH 10.5 (∼5,000 µmol kg^-1^ TA) and pH 9.0 (∼3,150 µmol kg^-1^ TA) for durations that reflect expected short-term exposure times during field OAE deployments (pH 10.5: 1, 5, 10 minutes; pH 9.0: 1, 15, 30 minutes). None of the treatment combinations resulted in mortality immediately after the initial exposure. Individuals were monitored for survival for 72 hours post-exposure (hpe), and only one treatment group (juveniles exposed to pH 10.5 for 10 minutes) showed a significant reduction in final survival; no other pH–duration combination showed increased mortality. Effects on the ability to initiate an escape response were more substantial. Adult females treated with pH 10.5 for 5 or 10 minutes showed a significant reduction in escape response immediately after exposure. In contrast, juveniles showed no immediate change in escape response following exposure to pH 10.5 or pH 9.0, although juveniles exposed to pH 10.5 for 10 minutes exhibited reduced escape response at 24 hpe. Using microrespirometry, we measured oxygen consumption following a 10-minute exposure to pH 10.5 and detected no effect on routine metabolic rate immediately post-exposure or at 12 hpe. Overall, our results suggest that *C. finmarchicus* is relatively tolerant to short-term exposures to very high pH and alkalinity. Future work should prioritize longer-term exposure under more moderate ocean OAE conditions.

## Introduction

Ocean Alkalinity Enhancement (OAE) is a marine carbon dioxide removal (mCDR) strategy that involves adding alkaline substances to the surface ocean to increase its capacity to absorb and store atmospheric CO_2_. In addition to climate change mitigation, OAE has the potential to increase seawater buffering capacity and counteract ocean acidification (Feng *et al*. 2017). OAE is therefore emerging as a popular CO_2_ removal strategy in voluntary carbon markets, with major corporations investing in its potential (Oschlies *et al*. 2023). Multiple field trials to assess the efficiency and practicality of this technology are currently underway or have been recently completed (Guo *et al*. 2025, Savoie *et al*. 2025, Subhas *et al*. 2025, Wang *et al*. 2025, Wynn-Edwards *et al*. 2025).

Despite this momentum, there is still considerable uncertainty about the potential for negative biological impacts (Iglesias-Rodríguez *et al*. 2023). The dispersal of concentrated forms of alkalinity into the surface ocean will rapidly increase pH and alkalinity at rates and magnitudes which greatly exceed levels of natural variability in most marine systems (Subhas *et al*. 2023). Aquatic animals within the near-field dispersal zone may experience acute stress due to the rapid rise in seawater pH and buffering capacity, as well as the corresponding decline in carbon dioxide partial pressure (*p*CO_2_). Depending on severity and duration of exposure, OAE conditions may disrupt essential physiological processes around acid-base homeostasis, ammonia excretion, gas exchange, and sensory function that may ultimately impact growth, reproduction, and survival (Ip and Chew 2010, Ashur *et al*. 2017, Fehsenfeld and Weihrauch 2017, Tresguerres *et al*. 2020).

Specifically, the response of marine animals to OAE conditions remains insufficiently characterized, which limits our ability to forecast ecological consequences for large-scale OAE deployments. Small zooplankton, such as copepods, may be disproportionately vulnerable to rapid changes in ocean chemistry. Copepods lack true gills and rely on cutaneous transport across body surfaces to exchange ions, gases, and metabolic waste products with the environment (Johnson *et al*. 2014). Their high surface-area-to-volume ratio means large external shifts in pH and alkalinity can more rapidly disturb internal pH and ion balance (McAllen *et al*. 1998). When energy reserves are limited, the added energetic costs of ion regulation may compromise growth, movement, and reproduction (Thor *et al*. 2022). Despite their strong swimming capabilities (Lenz *et al*. 2004), a copepods’ small size limits their ability to avoid large OAE plumes, resulting in prolonged exposures to potentially stressful conditions. To date, only a few experimental studies have investigated copepod responses to elevated pH and alkalinity, showing impacts to survival, swimming activity, feeding, and respiration rates (Hansen *et al*. 2017, Camatti *et al*. 2024, Bhaumik *et al*. 2025). However, many of the most severe impacts were observed after exposure to very high pH (>9.0) for long durations (>24 hours), conditions that may not be representative of most OAE scenarios.

For example, open-ocean OAE applications that use shipboard dispersal of alkaline agents into the turbulent ship wake produce surface plumes with extreme but short-lived spikes in pH and alkalinity (+1.2 pH units and +1200 µmol kg⁻¹ total alkalinity), on the order of seconds to minutes, before turbulent mixing and dilution quickly restores pH and alkalinity to a more moderate condition closer to the baseline environment (see Fig. 1; Chou 1996, Caserini *et al*. 2021, He and Tyka 2023, Subhas *et al*. 2025). Point-source discharges from land-based outfall pipes have shown similarly rapid dilution within the immediate plume area (Wynn-Edwards *et al*. 2025). In such addition/dilution scenarios, the exposure intensity is inversely correlated with the exposure time; a relatively small volume (containing fewer organisms) experiences extreme conditions for short durations, while a larger volume, containing more organisms, experience moderate OAE conditions for a longer duration (Fig. 1). Therefore, targeted research is urgently needed on copepod responses to the rapid pH fluctuations associated with open-ocean-style OAE, as such high-frequency pH changes themselves may be stressful (Clark and Gobler 2016). Furthermore, examining potential impacts on copepods is especially important because they often dominate marine zooplankton assemblages and form a key trophic link, transferring energy from phytoplankton to fish and other higher-level consumers (Turner 2004). Changes in copepod abundance or productivity can have outsized effects on marine food-web dynamics and stability, with broad implications for fisheries productivity (Alvarez-Fernandez *et al*. 2015).

**Fig. 1:**
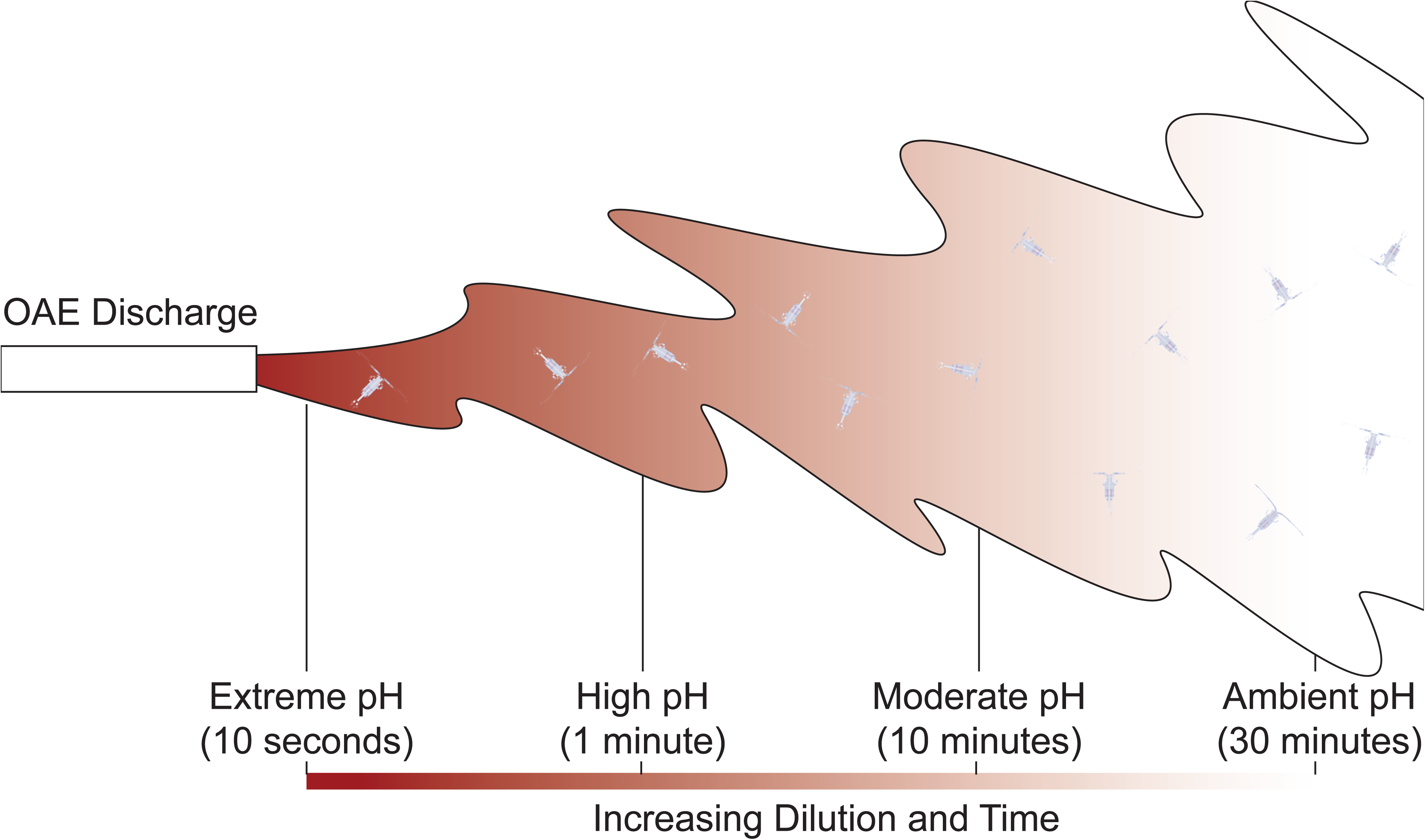
A conceptual depiction of a point-source alkalinity addition and its subsequent dilution. Elevated pH and alkalinity at the point of discharge dilutes with surrounding seawater over a timescale of seconds to minutes, creating a range of exposure intensities and durations. As the size of plume increases, it becomes more diluted but will potentially interact with a larger number of animals.

To address this knowledge gap, we evaluated the effects of extreme OAE conditions on *Calanus finmarchicus*, a dominant calanoid copepod in the North Atlantic (Dale *et al*. 2001). The species’ high abundance and lipid-rich composition make it a primary food source for many fish, seabirds, and marine mammals (Helenius *et al*. 2024). *C. finmarchicus* has been the focus of ocean acidification research investigating the effects of low pH (Mayor *et al*. 2007, Preziosi *et al*. 2017), but no published studies have tested the effects of elevated pH and alkalinity conditions expected during OAE in this particular species. In this study, we used NaOH as the alkalinity feedstock which may emerge as a preferred alkalinity source for widescale OAE due to its absence of toxic trace metals and high effectiveness in increasing alkalinity (Iglesias-Rodríguez *et al*. 2023). We focused on the acute, near-field phase of NaOH dispersal, testing two elevated pH treatments with associated carbonate chemistry changes across multiple exposure durations to determine how exposure intensity and duration mediate physiological impacts.

We evaluated three response traits: routine metabolic rate (RMR), escape response, and survival. RMR is defined as the metabolic rate supporting basic maintenance (Chabot *et al*. 2016). Under environmental stress, energetic costs of homeostatic maintenance can increase, resulting in higher RMR and a reduced pool of energy available for growth and other processes (Murray and Klinger 2022). In copepods, RMR may increase under OAE conditions if ion and acid–base regulation become more energetically costly, but may decline if compensation fails and disturbances in gas exchange, internal pH, or ion balance trigger metabolic depression (Thor *et al*. 2016). The copepod escape response is an important anti-predatory defense mechanism that involves a high-speed, coordinated swimming reaction to a stimulus (Kiørboe *et al*. 2010). The integration of neuromuscular processes underlying this response can be impaired by acute physiological stress. For example, calanoid copepods exposed to near-lethal concentrations of hydrogen peroxide, a common treatment against parasitic infestation in marine aquaculture, showed dramatic declines in escape performance (Escobar-Lux *et al*. 2019). Furthermore, exposure to extreme pH conditions may overwhelm compensatory mechanisms, leading to increased short-term mortality rates. We hypothesized that acute high-pH exposure would (i) increase short-term mortality, (ii) depress copepod escape responses, and (iii) alter metabolic rates, with the size of the effect increasing with the magnitude of the pH elevation and with the duration of exposure.

## Methods

### Field Collections and Laboratory Maintenance

Wild *C. finmarchicus* were collected twice from different locations in the Gulf of Maine. On 7 April 2025, copepods were sampled from Wilkinson Basin (42.86008° N, 69.86290° W) with vertical plankton tows using paired 200 µm mesh nets (mouth diameter 0.75 m), from a maximum depth of 250 m. A second collection occurred on 7 May 2025 in Cape Cod Bay (41.85181° N, 70.42978° W) using vertical tows with a paired bongo net (150 µm and 300 µm mesh; mouth diameter 0.75 m per ring) from a maximum depth of 25 m. After each tow, the bulk catch was diluted into 20-L containers fitted with battery-powered bubblers for gentle aeration. The containers were transported in large, insulated coolers to maintain near-ambient temperature to Woods Hole Oceanographic Institution.

The first collection yielded ∼300 adult females (copepodite stage 6, the final mature developmental stage) (Hansen *et al*. 2024). Adult females were maintained in 60-L circular polyethylene tubs with a slow flow-through of ambient laboratory filtered seawater (∼50 mL min⁻¹; ∼1.2 volume exchanges day⁻¹) sourced from Vineyard Sound [pH (total scale) 7.95; salinity 34 ppt; temperature 14–15 °C; total alkalinity (TA) 2170 µmol kg⁻¹]. The second collection yielded several thousand *C. finmarchicus* across multiple life stages, along with other copepod species (e.g., *Pseudocalanus* spp. and *Centropages typicus*). Among *C. finmarchicus*, late-stage copepodites (C4–C5; hereafter referred to as juveniles) were the most abundant and were selected for experiments. More than 1,000 juveniles were selected and distributed among four 60-L circular tubs supplied with ambient laboratory seawater [pH 7.95; salinity 34 ppt; temperature 16–18.5 °C; TA 2150 µmol kg⁻¹]. Gentle bubbling via air stones kept dissolved oxygen near saturation.

Each group of copepods was maintained in the laboratory for several weeks (adult females: 3 weeks; juveniles: 5 weeks) while experiments were conducted. Copepod holding tubs were checked daily for ammonia (API Ammonia Test Kit) and levels remained undetectable. The tubs were carefully siphoned every other day to remove waste. The photoperiod of the seawater laboratory was automatically adjusted to match the natural day:night cycle. Copepods were fed *ad libitum* daily rations of a mixed phytoplankton diet: *Thalassiosira weissflogii*, *Heterocapsa* spp., *Isochrysis galbana*, and *Rhodomonas salina*. The four algal cultures were grown on f/4 silicate media (Bigelow National Center for Marine Algae), kept in the exponential growth phase at 16 °C, and harvested daily.

### OAE Treatments

Copepods were exposed to two OAE treatments representative of near-field conditions that would follow an open-ocean dispersal of NaOH: an extreme treatment with a target pH of 10.5 (∼5,170 µmol kg^-1^ TA, +3,020 µmol kg^-1^ above ambient), and a moderate treatment with a target pH of 9.0 (∼3,180 µmol kg^-1^ TA, +1,030 µmol kg^-1^ above ambient). The extreme OAE treatment reflects the maximum pH and alkalinity conditions achievable in small-scale NaOH dosing trials before excessive precipitation of calcium carbonate and brucite and may represent the near-upper limit that organisms would experience immediately following NaOH dispersal in the field (Ringham *et al*. 2024). The moderate OAE treatment was selected to represent longer-duration conditions following NaOH dispersal in the surface ocean, while still exceeding the U.S. EPA’s recommended upper pH limit of 8.5 for ocean waters (U. S. Environmental Protection Agency 1986). A control treatment was paired with each trial using ambient laboratory seawater (∼7.95 pH, ∼2,150 µmol kg^-1^ TA).

Each pH treatment was applied at three exposure durations designed to match or conservatively exceed expected pH-specific exposure durations during OAE field trials (Table 1). Durations were informed by ship-wake models incorporating realistic dispersal rates, which suggest that pH increases of +1 to +1.5 are typically confined to minutes (<5 min) within the near-field plume, whereas more extreme pH values approaching 10.5 are expected to persist for seconds in the immediate vicinity of the alkalinity release (Caserini *et al*. 2021, Subhas *et al*. 2025).

**Table 1.** Summary of experimental trials on adult and juvenile *C. finmarchicus* assessing the effects of elevated pH and alkalinity on survival and escape response. Each trial included two target pH (total scale) treatments: a control group (pH 7.95) and an elevated pH treatment (pH 9.0 or 10.5) applied for three exposure durations (minutes). Reported here are experimental units (N), trial temperature (temp; °C), initial and final pH measured with a benchtop pH probe (converted from NIST to total scale with CO2SYS), mean total alkalinity from initial and final samples (TA; µmol kg^-1^), and mean dissolved inorganic carbon (DIC; µmol kg^-1^) from control exposure tubs. Analytical precision for TA and DIC is reported as ± standard deviation of three replicate measurements per seawater sample, drawn from an initial sample only (control pH) or from pooled initial and final samples (elevated pH/alkalinity treatments). Calculated pH values were derived with CO2SYS based on direct measurements of TA, DIC, temperature, and salinity to compare against the initial and final pH measurements made with the benchtop electrode.

| Trial | Life Stage | Target pH | Duration (minutes) | N | Temp (°C) | Initial pH | Final pH | Measured TA ($\mu\text{mol kg}^{-1}$) | Measured DIC ($\mu\text{mol kg}^{-1}$) | Calculated pH |
| --- | --- | --- | --- | --- | --- | --- | --- | --- | --- | --- |
| 1 | adult | 7.95 | ----- | 3 | 15.2 | 7.96 | 7.97 | 2149±0.72 | 1981±0.83 | 7.96 |
| | | 10.5 | 1,5,10 | 3 | 15.2 | 10.53 | 10.49 | 5012±2.62 | ----- | 10.49 |
| 2 | juvenile | 7.95 | ----- | 5 | 16.2 | 7.94 | 7.95 | 2166±1.83 | 1998±0.70 | 7.94 |
| | | 10.5 | 1,5,10 | 5 | 16.2 | 10.44 | 10.42 | 4925±5.20 | ----- | 10.39 |
| 3 | juvenile | 7.95 | ----- | 5 | 17.6 | 7.89 | 7.89 | 2138±2.95 | 2007±0.44 | 7.83 |
| | | 10.5 | 1,5,10 | 5 | 17.6 | 10.46 | 10.44 | 5322±3.22 | ----- | 10.52 |
| 4 | juvenile | 7.95 | ----- | 5 | 17.0 | 7.97 | 7.95 | 2146±3.22 | 1998±0.26 | 7.88 |
| | | 9.0 | 1,15,30 | 5 | 17.0 | 9.02 | 8.98 | 3169±1.75 | ----- | 9.00 |
| 5 | juvenile | 7.95 | ----- | 5 | 17.6 | 7.94 | 7.94 | 2148±1.46 | 2001±0.77 | 7.87 |
| | | 9.0 | 1,15,30 | 5 | 17.6 | 9.02 | 8.98 | 3172±2.22 | ----- | 8.99 |

NaOH-dosed seawater was prepared immediately before each trial in 4-L batches of filtered (to 1 µm) seawater. Seawater was gently stirred in HDPE beakers while a 50% NaOH solution was added in 10 µL increments to limit localized supersaturation and minimize precipitation. Target additions were 163 µL L^-1^ for the extreme OAE treatment (3.10 mM NaOH; ΔTA = +3,020 µmol kg^-1^) and 55 µL L^-1^ for the moderate OAE treatment (1.05 mM NaOH; ΔTA = +1,030 µmol kg^-1^).

### Experimental Design

OAE exposure experiments were conducted over five trials (Table 1). The limited number of adult female *C. finmarchicus* available from the first collection permitted just a single trial that tested pH 10.5. The second collection supported four trials with juvenile *C. finmarchicus*: two replicate trials testing pH 10.5 and two replicate trials testing pH 9.0 (Table 1). Each trial used two 40-L rectangular exposure tubs (24″ W × 18″ H × 6″ D), one assigned to the treatment pH (filled with 12 L of NaOH-dosed seawater) and one to the control (12 L of ambient seawater). Initial pH in exposure tubs was verified with a benchtop pH meter (Fisher Scientific Accumet AE150) calibrated with three NIST-traceable buffers.

Copepods were exposed to treatment conditions in custom floating vessels (0.5-L plastic cups with 150-µm mesh bottoms) that were designed to facilitate the rapid transfer of groups of small organisms between seawater treatments (Murray and Klinger 2022). These vessels served as the unit of experimental replication. Before each trial, healthy copepods were selected and distributed among replicate vessels for each treatment group (pH × duration combination) and the control group. Trial 1 used N = 3 vessels per group with 5 copepods per vessel. Trials 2–5 used N = 5 vessels per treatment group with 12–15 copepods per vessel. All replicate vessels were initially submerged in a water table with control seawater. To initiate the exposure, vessels assigned to the elevated-pH treatment were transferred to an exposure tub holding NaOH-dosed seawater for their designated exposure duration. Control vessels underwent a transfer to a control seawater tub to reproduce the effects of handling and air exposure. Final pH in the exposure tubs was rechecked at the end of the trial (trials lasted >4 h) with the benchtop pH meter, showing that elevated pH levels were relatively stable during the experiments and that declines due to CO₂ uptake were small (<0.05 units; Table 1).

### Seawater Carbonate Chemistry

Control seawater was sampled at the start of each trial into 60 mL borosilicate vials, sealed without headspace, poisoned with mercuric chloride (0.02 % final concentration), and stored in the dark at room temperature for direct measurements of TA (µmol kg^-1^) and dissolved inorganic carbon (DIC; µmol kg^-1^). NaOH-dosed seawater was sampled at the beginning and end of each trial and underwent additional processing to preserve elevated TA conditions (Schulz *et al*. 2023). Samples were acidified by gently bubbling CO₂ gas (99% purity) for five seconds. Acidification effectively prevented mineral precipitation, but this step precluded measurements of DIC. Temperature and salinity (via refractometer) were recorded at the time of sampling. TA was measured using an open-system Gran titration on 5 ml samples in triplicate, using a Metrohm 805 Dosimat and a robotic Titrosampler. DIC was measured on control samples in triplicate using an Apollo SciTech AS-C6L DIC analyzer equipped with a LI-COR 7815 CO_2_ detector. Measurements of TA and DIC were calibrated against seawater-certified reference materials (Batch 206; Dickson 2024). CO2SYS (v2) was used to calculate the full suite of carbonate chemistry parameters (pH, partial pressure of CO₂ (*p*CO₂), carbonate and bicarbonate concentrations, and saturation states for calcite and aragonite) (Pierrot *et al*. 2011). For the elevated pH/alkalinity treatments, we used DIC values measured from control seawater samples from that trial as the input for CO2SYS to calculate the initial carbonate chemistry of treatment seawater. The full suite of carbonate chemistry parameters for each sample (including respirometry trials) are reported in Table S1.

### Survival and Escape Response

After each exposure duration, elevated pH vessels were returned to the control seawater table. Each copepod was then gently poured into a large Petri dish and immediately examined under a dissecting microscope for two binary outcomes: survival (alive/dead) and, among survivors, escape response (successful/unsuccessful). Escape response was evaluated through tactile stimulation by gently prodding the copepod’s antennae with the tip of a transfer pipette. A successful escape was recorded if the copepod jumped more than one body length. If no jump occurred after three prodding attempts, spaced 5 seconds apart, the response was recorded as a failure. Replicate exposures were staggered to allow sufficient time to evaluate copepods before the next vessel required assessment, such that all trials were completed within 3-4 hours. Surviving copepods were returned to their original vessels and maintained in ambient seawater without food for 72 hours post-exposure (hpe). Survival and escape response were reassessed at 24 and 72 hpe.

### Metabolic Rate

RMR was measured using closed respirometry with a Loligo Microplate Respirometry System with AutoResp3 software and a 24-well plate (200 µL well size). The system was calibrated prior to each assay following manufacturer’s guidelines. Treatment effects on RMR were investigated in adult females and in juveniles in duplicate assays for each life stage. The day before the assay, copepods were separated from holding tubs and held overnight without food. For adult females, two treatments were tested: a control and pH 10.5 exposure for 10 minutes (n = 5 – 6 copepods per treatment, 12 blanks per assay). Juveniles were tested under three pH treatments: control, pH 10.5 (10-minute exposure immediately before the assay); and a pH 10.5 + 12 h treatment (10-minute exposure 12 h before the assay) (n = 6 copepods per treatment, 6 blanks per assay). Elevated pH exposures were completed as described above. Immediately post-exposure, copepods were transferred back to control seawater and immediately loaded into the microplate using a plastic transfer pipette with a cut tip. All microplate wells were overfilled with control pH seawater (0.4-µm filtered) with dissolved oxygen at ∼100% air saturation (a.s.). Empty wells were filled with the same experimental seawater to serve as blanks to measure background respiration rates. Once all copepods were loaded, plates were sealed with adhesive sealing film, covered with a silicone pad, and a weighted block to limit gas exchange, then placed on the SDR reader inside a dark, temperature-controlled incubator set to a constant 15 °C. Oxygen measurements were initiated immediately, with oxygen concentration (% air saturation) recorded every four seconds. Assays ran for 70-80 minutes, after which copepods were preserved in 70% ethanol and dried at 60 °C for 6 h to determine individual dry mass (DM, 0.001 mg) on a microbalance. Duplicate assays for each life stage were conducted consecutively on the same day.

Preliminary data handling included the inspection of oxygen traces for each well. To account for sensor equilibration after the microplate was placed in the incubator, we excluded the initial 10 minutes of each assay, based on the stabilization of oxygen readings in the blank wells. After this settling period, oxygen concentration in the blanks changed minimally over the remainder of each assay, declining by <2% a.s. in all cases. Oxygen traces were screened for anomalies, and for two samples, brief segments in the beginning of the trace consistent with microbubble release were trimmed. Data recorded below 40% a.s. were also trimmed to avoid the effects of potential oxygen limitation. After trimming, only samples with at least 10 minutes of continuous data were retained for analysis. The mean (±standard deviation) analyzed oxygen trace duration was shorter for adult females (43±19 minutes) than for juveniles (58±5 minutes). Final oxygen traces for analyzed samples are shown in Fig. S1 and Fig. S2.

### Statistical Analysis

All analyses were conducted in R (v4.0.2) using RStudio (v1.3). Statistical significance was set at α = 0.05. Survival and escape responses of adult females in Trial 1 were analyzed separately from juveniles Trials 2–5. For Trials 2–5, duplicate trials at the same pH were analyzed together as experimental replicates (pH 10.5: Trials 2 & 3; pH 9.0: Trials 4 & 5). Final survival (at 72 hpe) and escape response were modeled as binomial counts per replicate vessel using bias-reduced binomial generalized linear models (GLMs) with a logit link to mitigate small-sample bias and issues of near-complete separation (R package brglm2; Kosmidis 2024). Treatment was modeled as a fixed factor with four levels: pH 8.0 (control) and three elevated pH–duration levels (pH 10.5: 1, 5, and 10 minutes; pH 9.0: 1, 15, and 30 minutes). For analysis of juvenile traits that combined multiple trials, trial was included as a blocking factor. For escape response, models also included a treatment × evaluation time interaction to test whether treatment effects varied at 24 and 72 hpe following the initial exposure. Model summaries for GLMs are provided in Table S2 and S3. We obtained estimated marginal means (EMMs) for survival and escape response probabilities from the fitted models (Table S4 and S5) using the emmeans::emmeans() function (R package emmeans; Lenth 2018).

We used Wald χ² tests (emmeans::joint_tests()) to assess the significance of the main effect of treatment on survival (Table S6), and the effect of treatment and the treatment × evaluation time interaction on escape response (Table S7). For escape response models, the tests used cluster-robust covariance estimates (CR2) computed with clubSandwich::vcovCR() (clustered by replicate ID) to account for repeated measurements on the same replicates over the monitoring period (R package clubSandwhich; Pustejovsky 2017). Dunnett’s tests were then used to determine if survival in elevated pH treatments differed significantly from the control group (Table S8). For escape response, Dunnet’s tests were conducted within each evaluation time point (Table S9). Model diagnostics were evaluated using performance::check_model(), which indicated no major violations associated with overdispersion, residual patterns, or influential observations (R package performance; Lüdecke *et al*. 2021).

For respirometry analysis, oxygen concentrations were first converted from % a.s. to µmol L^-1^ using the respirometry::conv_o2 (R package respirometry; Birk 2016). Oxygen consumption rates were analyzed with a rolling (moving-window) linear regressions (Karlsson and Søreide 2023). Rates were computed over 2-minute windows that included a 1-minute overlap with the previous window and iterated until >1 minute of data remained. For each window, the mean rate of oxygen change of blank wells was subtracted from each copepod rate to obtain corrected rates that accounted for background respiration. Final oxygen consumption rates were calculated by trimming the upper and lower 5% of rolling-slope values to exclude extreme values, potential artifacts, and brief periods of elevated movement. We then calculated the mean of the remaining 90% of rolling-slope values and used it as the final oxygen consumption rate.

Rates were converted from µmol L^-1^ O_2_ h^-1^ to µmol O_2_ h^-1^ by scaling to well volume (200 µl). Oxygen consumption rates were not significantly correlated with dry mass in adult females (mean DM = 0.144±0.025 mg, Pearson’s r = 0.21, p = 0.350) or juveniles (mean DM = 0.043±0.007 mg, Pearson’s r = 0.32, p = 0.058). However, to facilitate size-standardized comparisons, all final RMR values were normalized to DM and reported as µmol O_2_ h^-1^ mg^-1^. Treatment effects on RMR were tested separately for adult females and juveniles using car::Anova() (R package car; Fox and Weisberg 2018). Models included treatment as a fixed effect and assay as a blocking factor to account for between-trial variation. Outliers were identified with performance::check_outliers() using a robust z-scores (outliers flagged at robust z-score > 3). When outliers were detected, the analysis was repeated with and without the flagged observations to evaluate the sensitivity of their removal on overall treatment effects.

## Results

### Survival

In Trial 1, exposure of adult females to pH 10.5 for 1, 5, and 10 minutes did not result in any immediate mortalities (Fig. 2A). By 24 and 72 hpe, additional mortalities occurred in the 5-and 10-minute pH 10.5 exposure groups, resulting in lower average final survival in those groups (Fig. 2A). However, Wald tests showed no overall effect of treatment on final survival (Fig. 2B, Table S6). Notably, the model produced wide 95% confidence intervals for survival EMMs, reflecting limited precision due to the modest number of copepods tested per treatment (n = 15) and relatively few mortality events (1–4 deaths per treatment).

**Fig. 2.**
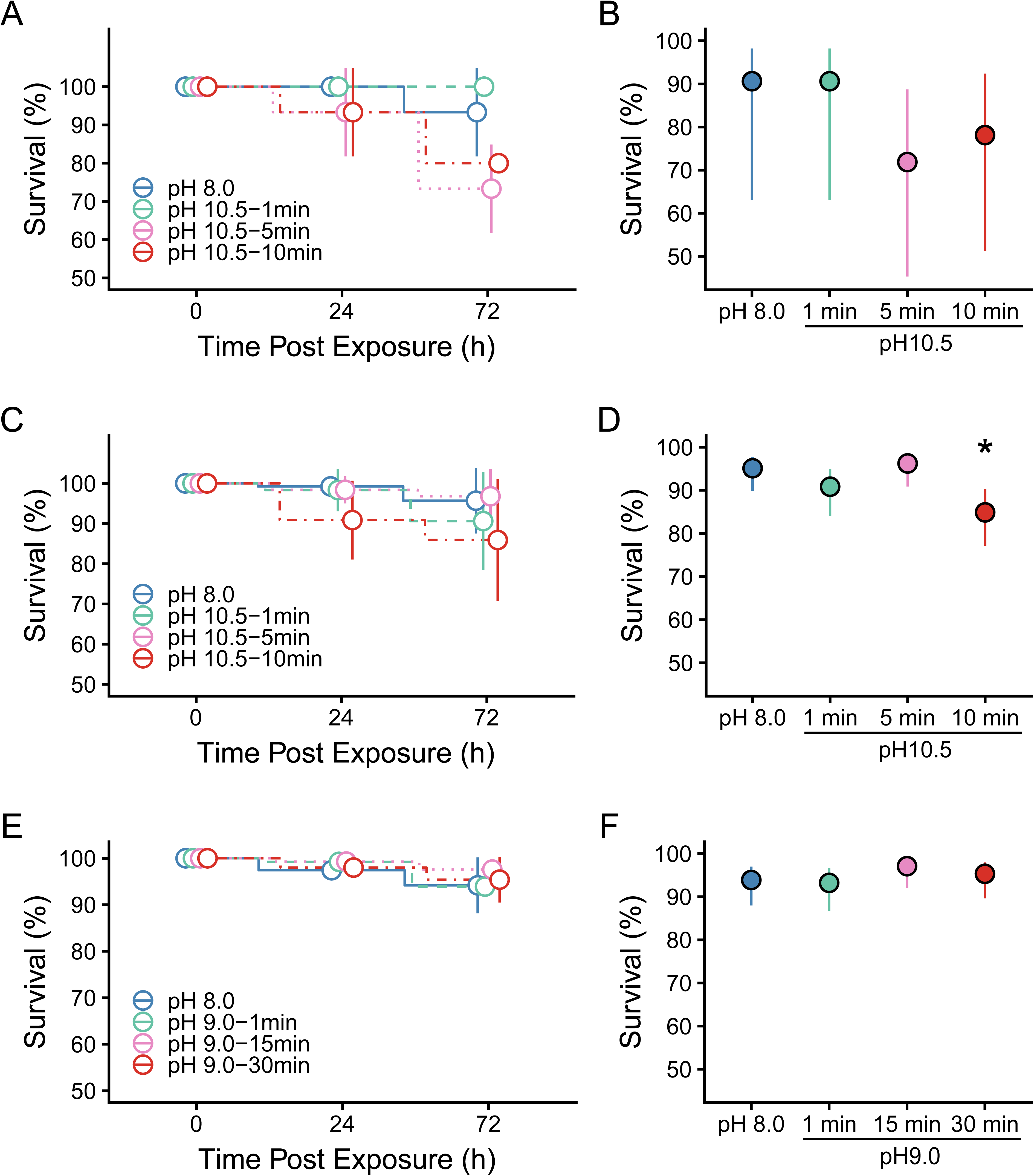
Survival of *C. finmarchicus* following exposure to elevated pH and alkalinity conditions in NaOH-dosed seawater. Results for adult females exposed to pH 10.5 (Trial 1; N = 3 vessels per treatment, n = 5 copepods per vessel) are shown in panels A and B, and results for stage juveniles exposed to pH 10.5 and 9.0 (Trials 2–5; N = 5 vessels per treatment, n = 12–15 copepods per vessel) are shown in panels C–F. Panels A, C, and E show survival over time post-exposure by treatment group. Open circles indicate the observed mean survival for each treatment at each time point, error bars show the standard deviation among replicate vessels, and dashed lines connect means between observation times. Panels B, D, and F show modeled final survival (72 h post-exposure) as estimated marginal means (EMMs) from a binomial generalized linear model. Treatment groups are labeled on the x-axis. Filled circles indicate EMMs by treatment, and error bars show 95% confidence intervals reflecting model-based uncertainty in final survival probability. An asterisk denotes treatments with survival probabilities that were significantly different from the control group (Dunnett’s test, p < 0.05).

In Trials 2 and 3, juvenile groups exposed to pH 10.5 for 1, 5, or 10 also showed no immediate mortality (Fig. 2C). Mean survival remained high at 24 hpe (>93% in all treatments; Fig. 2C). At 72 hpe, Wald test indicated a significant overall effect of treatment (Wald χ² = 11.526, *p =* 0.009, Table S6). The pH 10.5 group exposed for 10 minutes had significantly lower final survival (–10.7%) than the control group (Dunnet’s adjusted *p =* 0.026, Table S8), whereas survival in the 1– and 5-minute exposure groups did not differ from the control (Fig. 2D).

In Trials 4 and 5, juveniles showed no immediate mortality after exposure to pH 9.0 for 1, 15, and 30 minutes (Fig. 2E). Survival exceeded 97% across all groups at 24 hpe (Fig. 2E). At 72 hpe, final survival remained high across all treatment groups, with no significant differences between control and pH 9.0 treatments (Fig. 2F, Table S6).

### Escape Response

In Trial 1, the escape response of adult females was affected by significant treatment × evaluation time interaction (Wald χ^2^ = 29.436, *p* < 0.001, Table S7). Immediately after exposure, adult females exposed to pH 10.5 for 5 or 10 minutes had significantly lower escape response probabilities than controls (Dunnett-adjusted *p* = 0.005 for both comparisons; Fig. 3A, Table S9), but there was no effect in the group exposed to pH 10.5 for 1 minute. At 24 hpe, escape responses in pH 10.5 5-minute group remained significantly lower in the control (Dunnett-adjusted *p* = 0.031, Fig. 3A, Table S9), with no effects in the other two treatments. By 72 hpe, escape response probabilities did not differ between any pH 10.5 exposure group and the control (Fig. 3A).

**Fig. 3.**
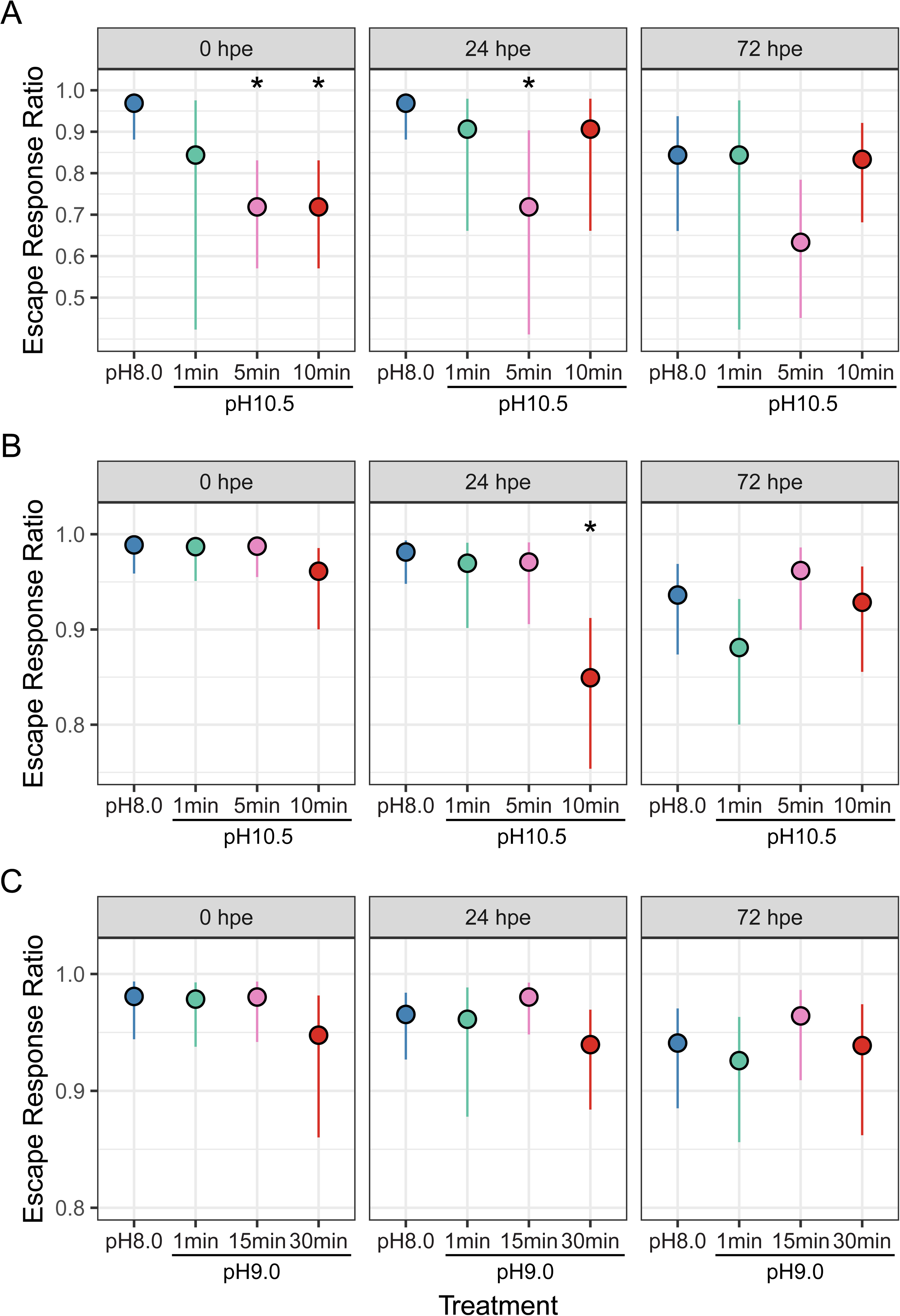
Estimated marginal means (EMMs) of escape response probability for *C. finmarchicus* following exposure to NaOH-dosed seawater. Panel A shows results for adult females tested at pH 10.5 (Trial 1; *N* = 3 vessels per treatment, *n* = 5 copepods per vessel), and panels B and C show results for juveniles tested at pH 10.5 and pH 9.0 (Trials 2–5; *N* = 5 vessels per treatment, *n* = 12-15 copepods per vessel). Panels from left to right show each observation time point (hours post-exposure, hpe) for each analysis group, treatments are labeled on the x-axis. Circles represent modeled EMMs of successful escape response ratios per treatment group. Error bars indicate 95% confidence intervals reflecting model-based uncertainty in escape response probability. An asterisk denotes treatments with escape response probabilities that were significantly different from the control group (Dunnett’s test, p < 0.05).

In Trials 2 and 3, the escape response of juveniles was also affected by a treatment × evaluation time (Wald χ^2^ = 17.352, *p* = 0.0081, Table S7) on the escape response of juveniles. In this case, there was no effect on escape response immediately post-exposure (Fig. 3B). At 24 hpe, the pH 10.5 10-minute group showed a significant reduction in successful escape response compared to the control group (Dunnet’s-adjusted *p* = 0.001, Table S9, Fig. 3B), whereas there was no effect in the pH 10.5 1– and 5-minute groups. By 72 hpe, there were no differences in escape responses between the control and any of the pH 10.5 groups (Fig. 3B).

In Trials 4 and 5, escape responses of stage juveniles exposed to pH 9.0 were not affected by treatment (Wald χ^2^ = 4.743, *p* = 0.192, Table S7) and or by a treatment × time post-exposure interaction (Wald χ^2^ = 2.514, *p* = 0.867, Table S7). Immediately post-exposure for 1, 15, or 30 minutes, escape response percentages were high (>95%) and similar across groups (Fig. 3C). Escape responses remained high at 24 and 72 hpe (Fig. 3C).

### Metabolic Rate

The RMR of adult females was not affected by a 10-minute exposure to pH 10.5 immediately prior to the measurement (ANOVA, F = 1.129, *p* = 0.302; Fig. 4A, Table S10). Mean RMR was 0.12 ± 0.05 and 0.14 ± 0.05 µmol O₂ h^-1^ mg^-1^ in the control and pH 10.5 groups, respectively. Across treatments, individual RMR varied from 0.06 to 0.22 µmol O₂ h^-1^ mg^-1^ (Fig. 4A).

**Fig. 4.**
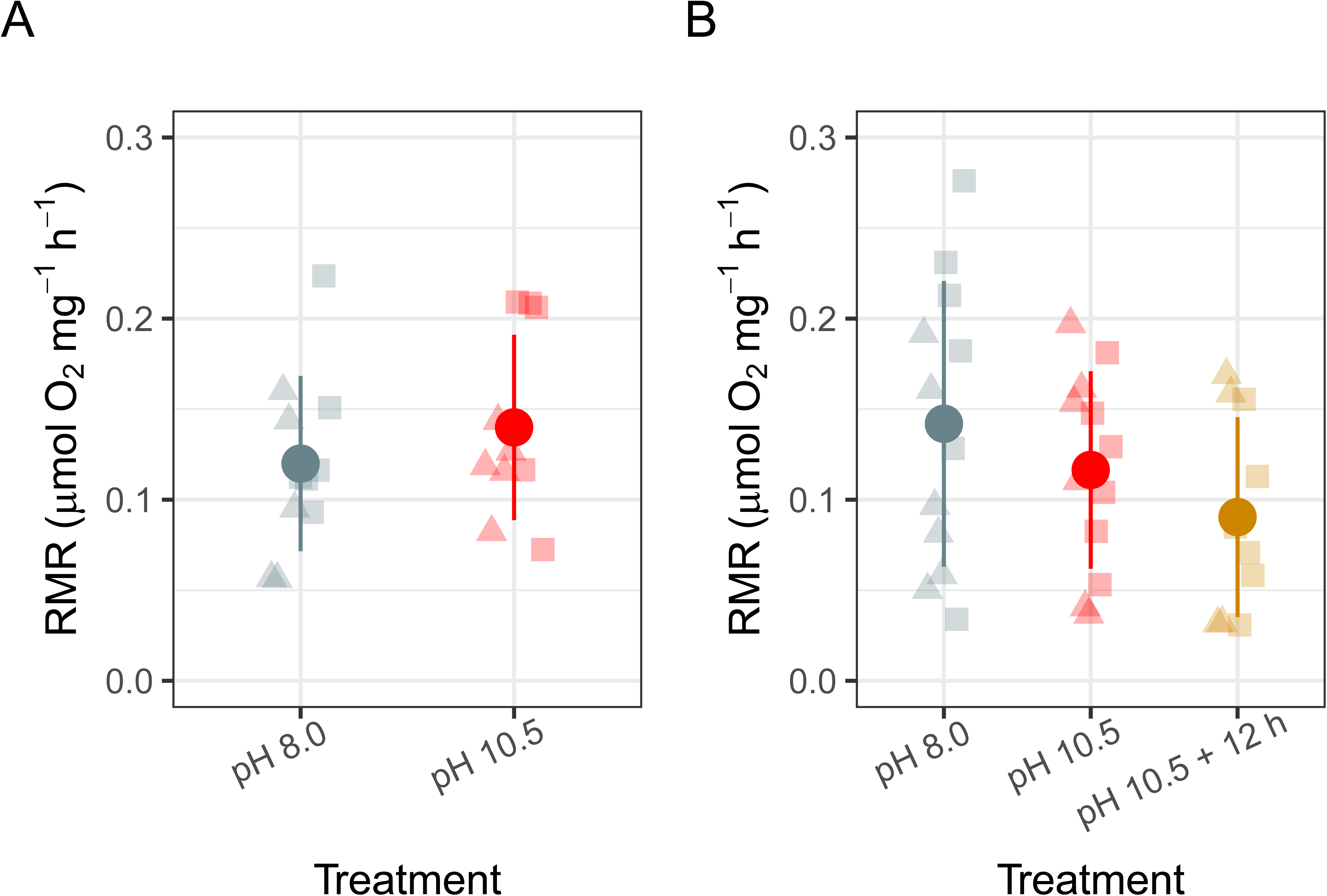
Effects of exposure to pH 10.5 on routine metabolic rates (RMR) of adult female (A) and juvenile (B) *C. finmarchicus* after statistical outliers were removed. For adult females, RMR was measured immediately post-exposure (pH 10.5). For juveniles, RMR was measured both immediately post-exposure (pH 10.5) and 12 hours post-exposure (pH 10.5 + 12 h). Large circles represent treatment means, and vertical bars indicate standard deviations. Individual RMR values are shown as small, semi-transparent triangles and squares, with different symbols corresponding to replicate RMR assays (n = 10–12 copepods per treatment, with 4–6 individuals per treatment in each replicate assay).

The RMR of juveniles was unaffected by treatment conditions (F = 0.471, *p* = 0.629; Table S10). However, two observations in the pH 10.5 + 12 h group were flagged outliers with relatively high RMRs of 0.68 and 0.41 µmol O₂ h^-1^ mg^-1^ (robust z-scores = 6.13 and 3.19). After excluding these observations, mean RMR was higher in the control group (0.14 ± 0.08 µmol O₂ h^-1^ mg^-1^) compared to the pH 10.5 (0.12 ± 0.05 µmol O₂ h^-1^ mg^-1^) and pH 10.5 + 12 h groups (0.09 ± 0.06 µmol O₂ h^-1^ mg^-1^) (Fig. 4B), but the overall effect of treatment remained non-significant (F = 1.883, *p* = 0.170, Table S10). Across individuals, RMR ranged from 0.03 to 0.28 µmol O₂ h^-1^ mg^-1^.

## Discussion

The identification of lethal thresholds for high pH is critically needed to appropriately scale field-based OAE experiments, and to ultimately regulate commercial applications. Therefore, there is an urgent need to design experimental studies that use specifically tailored exposure regimes to understand the effects of OAE in open-ocean settings. This study investigated how exposure to elevated pH consistent with near-field OAE plumes affects the keystone copepod *C. finmarchicus*, focusing on adult females and late-stage juveniles (copepodite stages 4 and 5). We implemented short-term exposure regimes at extreme (pH 10.5 for 1, 5, or 10 minute) and moderate (pH 9.0 for 1, 15, or 30 minute) OAE conditions to approximate the near-field environment following NaOH dispersal into the surface ocean. We hypothesized that these exposures would reduce short-term survival, but we found limited evidence of increased mortality. Juveniles exposed to pH 9 showed no effects on final survival evaluated at 72 hpe. At pH 10.5, a significant effect on survival was observed in juveniles exposed for 10 minutes, which showed a 10.7% reduction in final survival (at 72 hpe) compared to controls. There were no survival effects in the 1– and 5-minute exposure groups. Adult females showed a similar, albeit non-significant trend, with lower average final survival following pH 10.5 exposure for 5 and 10 minutes; however, these model estimates were less precise given the modest sample size.

Studies evaluating the lethality of elevated pH in copepods are scarce. One study tested the lethal effects of elevated pH in adults of six copepod species, finding LC_50_ values (the pH at which 50% of individuals died after a continuous 72-hour exposure) that ranged from 8.39 to 9.51 across taxa. The species with the most oceanic distribution, *Oithona similis*, showed the lowest LC_50_ (highest sensitivity) and was the only species to show mortality within a 24 h exposure. Whereas the most estuarine species, *Eurytemora affinis*, exhibited the highest tolerance, indicating that natural exposure to pH variability typical of coastal systems may contribute to a wider range of pH tolerance (Baumann 2019). Another study focused on *A. tonsa* reported dose– and time-dependent effects of high pH exposure on survival (Camatti *et al*. 2024). Copepods treated at pH 9 and 10 showed no change in survival after 24 hours of continuous exposure but exhibited moderate increases in mortality after 48 hours. In contrast, at pH 11, increased mortality was observed within 5 minutes, and complete morality was observed at pH 12 within minutes of initial exposure (Camatti *et al*. 2024). Negative effects were also observed in the marine copepod *Temora longicornis*, where a four-day exposure to pH 9.2 led to increased mortality (Bhaumik *et al*. 2025).

The emerging patterns suggest that lethal effects of elevated pH exposure are time-dependent, and aside from highest tested levels (pH ≥ 10.5), increased mortality occurs only after relatively long (e.g., 24 – 72 h) exposure. Most previous studies were not designed to assess lethal effects of short exposures of less than an hour. However, short exposures to very high pH levels may better reflect the conditions produced by OAE applications in the field. For example, in modeled ship-based OAE scenarios, alkaline material released into a turbulent ship wake is diluted by several orders of magnitude within minutes, and when combined with normal mixing in the ocean, organisms entrained in the near-field plume are likely to be exposed to the highest potential pH conditions (> 9.5) for seconds, up to a few minutes (Fig. 1; Caserini *et al*. 2021, Subhas *et al*. 2025). In our experiments, reduced survival was observed only after a 10-minute exposure to the extreme pH 10.5, suggesting this duration at pH 10.5 may represent a sensitivity threshold, with longer exposures potentially increasing short-term mortality rates. Additionally, because mortalities in this group continued to accumulate through 24 and 72 hpe (Fig. 2C), it may be necessary to evaluate longer-term post-exposure survival following extreme pH exposure, as lethal effects may accrue several days after the initial exposure.

We found that exposure to pH 10.5, but not pH 9.0, affected the copepods ability to initiate escape response, an important anti-predator behavior in which individuals detect the close-range hydrodynamic disturbances of potential predators and initiate a high-speed escape jump (Kiørboe *et al*. 2010). When assessed immediately after exposure, adult females exposed to pH 10.5 for 5 or 10 minutes showed ∼25% reduction in the initiation of escape responses. This difference was no longer detected in the 10-minute group by 24 hpe and was resolved in the 5-minute group by 72 hpe. In juveniles, there was no effect immediately following the exposures to pH 10.5 for any duration. By 24 h post-exposure, the pH 10.5 group exposed for 10 minutes showed a modest (∼8%) but significant reduction in escape response. The effect was no longer present by 72 hpe. Importantly, escape response was scored only among survivors at each timepoint. Thus, additional mortality in the pH 10.5 groups at 24 and 72 hpe may have influenced observed escape-response patterns over the monitoring period if non-responders died before the next assessment. This also suggests that failure to mount an escape response may predict forthcoming mortality.

A similar methodological approach was used to evaluate the escape activity of *A. tonsa*, where five minute exposures to pH 9 and 10 did not alter escape response frequencies, but exposure to pH 11 for five minutes significantly decreased the rate of escape responses, an effect that was further amplified after three hours of exposure (Camatti *et al*. 2024). Together, these data suggest that short-term exposure to extreme pH levels may compromise components of the neuromuscular processes underlying the copepod escape response. However, in this study, effects on *C. finmarchicus* were not consistent across 10.5 treatment groups and life stages (i.e., juveniles showed no immediate effects after 10.5 exposures). Furthermore, there were no effects observed in groups exposed to the more moderate pH 9.0 treatment. We note that the escape response metric evaluated in the present study was relatively coarse, capturing only whether individuals initiated a response or not. Our approach did not cover other key aspects of the behavior, such as distance threshold for inducing a response, escape swimming speeds, or jump kinematics, all of which are potentially sensitive to environmental stress and could be influenced by elevated pH (Kiørboe *et al*. 2010)

Using microplate respirometry, we found that a 10-minute exposure to pH 10.5 did not significantly affect RMR in either adult females or juveniles. We also tested for a delayed response in juveniles by measuring RMR 12 hours after the initial exposure and likewise found no effect. These findings do not support our hypothesis that RMR would show substantial changes following acute exposure to extreme pH. It is possible that, despite the extreme pH level, the brief exposure did not produce a measurable disturbance in internal acid–base or ion balance. Or, that the exposure duration was insufficient to elicit a detectable compensatory response before the copepod was returned to ambient seawater. For instance, in the calanoid copepod *T. longicornis*, a longer-term (four-day) exposure to moderately elevated pH conditions (pH 8.75–9.2) resulted in significant declines in respiration rate (Bhaumik *et al*. 2025). Furthermore, chronic exposure to low pH under ocean acidification scenarios resulted in increased metabolic rate and energetic turnover, likely due to elevated homeostatic maintenance costs (Thor *et al*. 2016, Thor *et al*. 2022, Espinel-Velasco *et al*. 2023). Thus, longer-term exposures, even under moderate pH and OAE conditions, may represent a more substantial physiological stressor than short-term exposure to extreme levels. Such chronic exposures may result in altered internal pH balance, limitations on gas exchange, or impaired elimination of nitrogenous waste, leading to metabolic dysfunction. However, in the case of *T. longicornis*, the high-pH effect on oxygen consumption was reversed when copepods fed on higher-quality prey (Bhaumik *et al*. 2025), indicating the potential for nutrition-related compensatory mechanisms, as has been observed in fish larvae and copepods under the effects of ocean acidification (Sswat *et al*. 2018, Meyers *et al*. 2019).

The mean mass-corrected RMR values reported here were very similar between adult females and juveniles, and were also comparable to previously reported mass-corrected oxygen consumption rates for *C. finmarchicus* collected during the spring season and grown under a similar temperature regime (Ingvarsdóttir *et al*. 1999). However, substantial within-group variance was evident, including in the control groups, with metabolic rates differing by several fold among individuals within the same treatment. Similar levels of inter-individual variability in metabolic rate measured by microplate systems have been reported for *C. finmarchicus* and other *Calanus* spp. (Karlsson and Søreide 2023), suggesting that a combination of true individual differences and uncontrolled factors (e.g., movement during measurement) can contribute to high variance in these assays. Furthermore, the 200-µL well size of the microrespirometry plate used in this study was well suited for juveniles but may have been slightly undersized for adult females, resulting in relatively short oxygen trace durations (<30 minutes) for some samples.

### Summary

Our results indicate that adult female and late stage copepodite *C. finmarchicus* are relatively tolerant of short-term exposure to elevated pH and alkalinity, with limited effects on survival, metabolic rate, and escape response. Exposure to pH 9.0 for up to 30 minutes produced no detectable effects on survival or escape response. At pH 10.5, RMR was unaffected. However, juveniles exposed to pH 10.5 for 10 minutes showed a small but significant reduction in 72-hour survival, whereas 1– and 5-minute exposures had no effect, suggesting a time-dependent sensitivity threshold at this extreme pH. Effects on escape response at pH 10.5 after 5– and 10-minute exposures were modest overall but warrant further study. Critically, the relative tolerance of this keystone copepod under extreme OAE conditions expected following ship-wake dispersal of NaOH into surface waters suggests that the short exposure durations anticipated in the field are unlikely to be lethal and are likely to have limited carryover effects on growth and fitness. Nevertheless, because some adverse effects emerged after 5– and 10-minute exposure at pH 10.5, dispersal protocols should aim to minimize organismal exposures to very high pH to avoid direct impacts on copepod survival and swimming performance.

A full understanding of broader food-web effects will require pairing these acute responses with measurements of more subtle sublethal effects that may be affected by extreme pH conditions. Additional sublethal metrics may include additional measurements of escape response dynamics, physiological stress markers (e.g., oxidative stress, heat shock proteins, metabolic enzymes), and long-term reproductive output, are promising areas for future research. In addition, it would be valuable to study the long-term, multi-day effects of chronic exposure to the moderately elevated pH (<9.0) and alkalinity levels (<+1000 µmol kg⁻¹) that characterize the steady-state surface ocean conditions following large-scale alkalinity dispersals.

## Author contributions

Conceptualization [C.S.M., A.S., J.R., L.M., N.A], Data curation [C.S.M.], Formal analysis [C.S.M.], Funding acquisition [A.S., J.R., A.W., K.C., H.H.K, A.M., D.M.], Investigation [C.S.M.], Methodology [C.S.M., N.A, L.M], Validation [C.S.M, A.S., J.R., L.M., N.A.], Visualization [C.S.M.], Writing–original draft [C.S.M.], Writing–review & editing [C.S.M.,L.M., N.A., A.S., J.R., A.W., K.C., H.H.K, A.M., D.M.].

## Data Availability

The survival, escape response, and respirometry data underlying this article are available in the Dryad repository (DOI: 10.5061/dryad.s7h44j1p1; reviewer share link: http://datadryad.org/share/LINK_NOT_FOR_PUBLICATION/bljGZ65L2Xe6SVZskJEexIgXnS7T5gM-soUku8hyKt4).

## Supplementary Material

The following supplementary material is available at ICESJMS online: Table S1: Carbonate chemistry measurements; Table S2: GLM summary for survival; Table S3: GLM summary for escape response; Table S4: EMMs of survival probabilities; Table S5: EMMs of escape-response probabilities; Table S6: Wald tests for survival; Table S7: Wald tests for escape response; Table S8: Dunnett’s tests for survival; Table S9: Dunnett’s tests for escape response; Table S10: ANOVA results for RMR, Fig. S1: oxygen traces from adult female respirometry; Fig. S2: oxygen traces from juvenile respirometry.

## Acknowledgements

We gratefully acknowledge Dr. Ann Tarrant and the crew of the R/V Gulf Challenger for facilitating the first collection of wild copepods on Wilkinson Basin. We thank Matt Hayden, Kate Morkeski, and the crew of the R/V Tioga for facilitating the second collection of copepods in Cape Cod Bay and to Chloe Dean for help with algal food culturing. We especially thank Phil Alatalo for support and guidance with copepod taxonomic identification, algal cultures, and general copepod rearing.

