## Supplemental Material for "The effects of elevated seawater pH and total alkalinity following dosing of sodium hydroxide in *Calanus finmarchicus*"

**Running head:** Effects of high pH in *Calanus finmarchicus*

**Keywords**: OAE, biological impacts, survival, escape response, routine metabolic rate, mCDR, copepod

### Supplemental Material

**Table S1. Carbon chemistry measurements**. Seawater samples were collected from each survival/escape response and respirometry (MO2) trial. MO2_1 corresponds to the trial on adult females and MO2_2 on juveniles. Target pH values are reported on the total scale. Timing indicates whether the sample was taken at the beginning (initial) or end (final) of the trial. Measured pH values (NIST scale) from a benchtop meter are reported for each sample (at the point of sampling), along with sample temperature (°C). All samples were analyzed for total alkalinity (TA; μmol $kg^{-1}$). Dissolved inorganic carbon (DIC; μmol $kg^{-1}$) was measured for control (target pH 7.95) samples only, because elevated-alkalinity samples were preserved via $CO_2$ bubbling. Standard deviations (SD) for TA and DIC are based on triplicate measurements for each seawater sample. CO2SYS was then used to calculate pH (total scale), $pCO_2$ (μatm), bicarbonate concentration ($HCO_3^-$; μmol $kg^{-1}$), carbonate concentration ($CO_3^{2-}$; μmol $kg^{-1}$), and the saturation states of calcite and aragonite using measured TA, DIC (from control samples as initial conditions), temperature, and salinity (34 ppt).

| Trial | Target pH (total) | Timing | Measured pH (NIST) | Sample temp | TA | TA sd | DIC | DIC sd | pH CO2SYS | $pCO_2$ CO2SYS | $HCO_3^-$ CO2SYS | $CO_3^{2-}$ CO2SYS | Cal CO2SYS | Arg CO2SYS |
|---|---|---|---|---|---|---|---|---|---|---|---|---|---|---|
| 1 | 7.95 | initial | 8.08 | 15.2 | 2149.67 | 0.56 | 1982.98 | 1.04 | 7.96 | 475.64 | 1842.44 | 122.80 | 2.95 | 1.89 |
| 1 | 7.95 | final | 8.09 | 15.2 | 2149.15 | 0.88 | 1978.58 | 0.62 | 7.97 | 463.82 | 1836.20 | 125.07 | 3.00 | 1.93 |
| 1 | 10.5 | initial | 10.65 | 15.2 | 5091.64 | 0.37 | | | 10.53 | 0.05 | 75.96 | 1904.04 | 45.69 | 29.31 |
| 1 | 10.5 | final | 10.61 | 15.2 | 4931.76 | 4.87 | | | 10.45 | 0.08 | 91.12 | 1887.46 | 45.30 | 29.05 |
| MO2_1 | 7.95 | initial | 8.11 | 15 | 2155.58 | 2.16 | 1972.29 | 0.88 | 8 | 427.02 | 1823.49 | 132.77 | 3.19 | 2.04 |
| MO2_1 | 10.5 | initial | 10.67 | 15 | 5165.37 | 4.27 | | | 10.58 | 0.04 | 68.26 | 1904.03 | 45.69 | 29.29 |
| 2 | 7.95 | initial | 8.06 | 16.2 | 2166.04 | 1.83 | 1998.28 | 0.70 | 7.94 | 498.78 | 1855.91 | 124.31 | 2.98 | 1.92 |
| 2 | 10.5 | initial | 10.56 | 16.2 | 4929.41 | 4.45 | | | 10.39 | 0.10 | 101.68 | 1896.60 | 45.54 | 29.28 |
| 2 | 10.5 | final | 10.54 | 16.2 | 4920.85 | 5.95 | | | 10.38 | 0.10 | 102.80 | 1895.48 | 45.52 | 29.26 |
| MO2_2 | 7.95 | initial | 8.03 | 15 | 2175.92 | 2.65 | 2025.88 | 0.93 | 7.92 | 536.66 | 1892.01 | 113.73 | 2.73 | 1.75 |
| MO2_2 | 10.5 | initial | 10.69 | 15 | 5253.68 | 4.87 | | | 10.57 | 0.04 | 71.32 | 1954.55 | 46.90 | 30.07 |
| 3 | 7.95 | initial | 8.01 | 17.6 | 2138.92 | 2.95 | 2007.42 | 0.44 | 7.83 | 657.69 | 1881.42 | 103.14 | 2.48 | 1.60 |
| 3 | 10.5 | initial | 10.58 | 17.6 | 5259.56 | 2.82 | | | 10.49 | 0.06 | 77.51 | 1929.91 | 46.39 | 29.92 |
| 3 | 10.5 | initial | 10.58 | 17.6 | 5357.29 | 4.21 | | | 10.53 | 0.05 | 70.84 | 1936.57 | 46.55 | 30.02 |
| 3 | 10.5 | final | 10.56 | 17.6 | 5314.05 | 3.11 | | | 10.51 | 0.05 | 73.66 | 1933.76 | 46.48 | 29.98 |
| 3 | 10.5 | final | 10.56 | 17.6 | 5356.22 | 2.73 | | | 10.53 | 0.05 | 70.91 | 1936.51 | 46.55 | 30.02 |
| 4 | 7.95 | initial | 8.09 | 17 | 2146.21 | 3.22 | 1998.19 | 0.26 | 7.88 | 576.94 | 1865.27 | 112.52 | 2.70 | 1.74 |

### Supplemental Material

| | | | | | | | | | | | | | | |
|---|---|---|---|---|---|---|---|---|---|---|---|---|---|---|
| 4 | 9 | initial | 9.02 | 17 | 3175.34 | 1.53 | | | 9.00 | 26.44 | 1116.94 | 880.31 | 21.15 | 13.62 |
| 4 | 9 | final | 8.98 | 17 | 3161.71 | 1.97 | | | 8.99 | 27.30 | 1127.89 | 869.34 | 20.89 | 13.45 |
| 5 | 7.95 | initial | 8.06 | 17.6 | 2143.45 | 1.90 | 2026.33 | 0.25 | 7.79 | 730.27 | 1905.65 | 95.30 | 2.29 | 1.48 |
| 5 | 9 | initial | 9.15 | 17.6 | 3171.11 | 3.83 | | | 8.96 | 30.58 | 1168.94 | 856.33 | 20.58 | 13.28 |
| 5 | 9 | final | 9.11 | 17.6 | 3173.62 | 0.60 | | | 9.01 | 24.91 | 1078.79 | 895.23 | 21.52 | 13.88 |

#### Supplemental Material

**Table S2 - General linear models (GLMs) of final survival.** Summaries of bias-reduced binomial GLMs with a logit link (R package brglm2) testing the effects of treatment (elevated pH–duration combinations) on final survival at 72 h post-exposure. Trial was included as blocking factor where necessary. Final survival was modeled as a two-column binomial response, representing the number surviving and dead from the initial number exposed. Results are shown for the three analysis groups.

| **Analysis Group** | **Coefficients** | **Estimate** | **Std. Error** | **Z value** | ***p*** |
|---|---|---|---|---|---|
| Adult Females (Trial 1) | Intercept | 2.269 | 0.886 | 2.561 | 0.0104 |
| | pH10.5-1min | 4.043e-16 | 1.253 | 0.000 | 1.0000 |
| | pH10.5-5min | -1.330 | 1.056 | -1.260 | 0.2076 |
| | pH10.5-10min | -0.9957 | 1.084 | -0.919 | 0.3583 |
| **Analysis Group** | **Coefficients** | **Estimate** | **Std. Error** | **Z value** | ***p*** |
| Juveniles (Trials 2-3**)** | Intercept | 2.534 | 0.654 | 3.876 | 0.0001 |
| | pH10.5-1min | -0.677 | 0.516 | -1.313 | 0.1892 |
| | pH10.5-5min | 0.270 | 0.626 | 0.432 | 0.6657 |
| | pH10.5-10min | -1.245 | 0.478 | -2.604 | 0.0092 |
| | Trial | 0.146 | 0.169 | 0.860 | 0.3898 |
| **Analysis Group** | **Coefficients** | **Estimate** | **Std. Error** | **Z value** | ***p*** |
| Juveniles (Trials 4-5**)** | Intercept | 2.866 | 0.967 | 2.963 | 0.0030 |
| | pH9.0-1min | -0.111 | 0.532 | -0.208 | 0.8353 |
| | pH9.0-15min | 0.776 | 0.657 | 1.181 | 0.2376 |
| | pH9.0-30min | 0.283 | 0.574 | 0.492 | 0.6224 |
| | Trial | -0.034 | 0.212 | -0.160 | 0.8733 |

#### Supplemental Material

**Table S3. General linear models (GLMs) of escape response**. Summaries of bias-reduced binomial GLMs with a logit link (R package brglm2) testing the interactive effects of treatment (elevated pH–duration combinations) and evaluation time (0-, 24-, and 72-hours post-exposure; hpe) on escape response. Escape response was modeled as a two-column binomial response, representing the number of individuals exhibiting an escape response and the number not exhibiting an escape response out of the total number of living copepods. Results are shown for the three analysis groups.

| Analysis Group | Coefficients | Estimate | Std. Error | Z value | *p* |
|---|---|---|---|---|---|
| Adult Females (Trial 1) | Intercept | 3.43 | 1.48 | 2.314 | 0.0207 |
| | pH10.5-1min | -1.75 | 1.65 | -1.062 | 0.2882 |
| | pH10.5-5min | -2.50 | 1.59 | -1.568 | 0.1168 |
| | pH10.5-10min | -2.50 | 1.59 | -1.568 | 0.1168 |
| | time_check-24hpe | 1.22e-15 | 2.10 | 0 | 1 |
| | time_check-72hpe | -1.75 | 1.65 | -1.062 | 0.2882 |
| | pH10.5-1min:time_check-24hpe | 0.58 | 2.39 | 0.244 | 0.8072 |
| | pH10.5-5min:time_check-24hpe | 1.58e-12 | 2.25 | 0 | 1 |
| | pH10.5-10min:time_check-24hpe | 1.33 | 2.35 | 0.566 | 0.5712 |
| | pH10.5-1min:time_check-72hpe | 1.75 | 1.93 | 0.906 | 0.3648 |
| | pH10.5-5min:time_check-72hpe | 1.36 | 1.83 | 0.741 | 0.4585 |
| | pH10.5-10min:time_check-72hpe | 2.42 | 1.89 | 1.283 | 0.1994 |
| **Analysis Group** | **Coefficients** | **Estimate** | **Std. Error** | **Z value** | ***p*** |
| Juveniles (Trials 2-3**)** | Intercept | 4.367 | 0.832 | 5.250 | 0.0000 |
| | pH10.5-1min | -0.155 | 1.162 | -0.134 | 0.8937 |
| | pH10.5-5min | -0.113 | 1.162 | -0.097 | 0.9226 |
| | pH10.5-10min | -1.272 | 0.952 | -1.336 | 0.1814 |
| | time_check-24hpe | -0.518 | 1.040 | -0.498 | 0.6183 |
| | time_check-72hpe | -1.796 | 0.894 | -2.008 | 0.0446 |
| | trial | 0.230 | 0.251 | 0.916 | 0.3598 |
| | pH10.5-1min:time_check-24hpe | -0.347 | 1.433 | -0.242 | 0.8089 |
| | pH10.5-5min:time_check-24hpe | -0.346 | 1.433 | -0.241 | 0.8092 |

| | | | | | |
|---|---|---|---|---|---|
| | pH10.5-10min:time_check-24hpe | -0.962 | 1.175 | -0.819 | 0.4128 |
| | pH10.5-1min:time_check-72hpe | -0.528 | 1.249 | -0.423 | 0.6725 |
| | pH10.5-5min:time_check-72hpe | 0.655 | 1.307 | 0.501 | 0.6162 |
| | pH10.5-10min:time_check-72hpe | 1.151 | 1.084 | 1.061 | 0.2886 |
| **Analysis Group** | **Coefficients** | **Estimate** | **Std. Error** | **Z value** | ***p*** |
| Juveniles (Trials 4-5**)** | Intercept | 4.148 | 0.667 | 6.220 | 4.99E-10 |
| | pH9.0-1min | -0.115 | 0.905 | -0.127 | 0.8990 |
| | pH9.0-15min | -0.026 | 0.904 | -0.029 | 0.9770 |
| | pH9.0-30min | -1.037 | 0.756 | -1.371 | 0.1700 |
| | time_check-24hpe | -0.605 | 0.800 | -0.756 | 0.4500 |
| | time_check-72hpe | -1.165 | 0.743 | -1.569 | 0.1170 |
| | trial | -0.430 | 0.279 | -1.542 | 0.1230 |
| | pH9.0-1min:time_check-24hpe | -0.002 | 1.132 | -0.002 | 0.9990 |
| | pH9.0-15min:time_check-24hpe | 0.605 | 1.207 | 0.501 | 0.6160 |
| | pH9.0-30min:time_check-24hpe | 0.453 | 0.973 | 0.465 | 0.6420 |
| | pH9.0-1min:time_check-72hpe | -0.128 | 1.044 | -0.122 | 0.9030 |
| | pH9.0-15min:time_check-72hpe | 0.550 | 1.092 | 0.504 | 0.6140 |
| | pH9.0-30min:time_check-72hpe | 0.999 | 0.926 | 1.078 | 0.2810 |

#### Supplemental Material

**Table S4. Estimated marginal means (EMMs) of final survival.** EMM survival probabilities (± standard errors) obtained from fitted GLMs for each pH-duration treatment combination. Survival probabilities are reported on the response scale. LCL and UCL denote the lower and upper 95% confidence limits. Results are shown for the three analysis groups.

| Analysis Group | Treatment | Probability | Std. Error | LCL | UCL |
|---|---|---|---|---|---|
| Adult Females (Trial 1) | pH 8.0 (control) | 0.906 | 0.075 | 0.630 | 0.982 |
| | pH10.5-1min | 0.906 | 0.075 | 0.630 | 0.982 |
| | pH10.5-5min | 0.719 | 0.116 | 0.453 | 0.887 |
| | pH10.5-10min | 0.781 | 0.107 | 0.512 | 0.924 |
| **Analysis Group** | **Treatment** | **Probability** | **Std. Error** | **LCL** | **UCL** |
| Juveniles (Trials 2-3) | pH 8.0 (control) | 0.951 | 0.019 | 0.899 | 0.977 |
| | pH10.5-1min | 0.908 | 0.027 | 0.840 | 0.949 |
| | pH10.5-5min | 0.962 | 0.017 | 0.909 | 0.985 |
| | pH10.5-10min | 0.849 | 0.033 | 0.771 | 0.903 |
| **Analysis Group** | **Treatment** | **Probability** | **Std. Error** | **LCL** | **UCL** |
| Juveniles (Trials 4-5) | pH8.0 (control) | 0.939 | 0.022 | 0.880 | 0.970 |
| | pH9.0-1min | 0.932 | 0.024 | 0.868 | 0.966 |
| | pH9.0-15min | 0.971 | 0.015 | 0.920 | 0.990 |
| | pH9.0-30min | 0.953 | 0.020 | 0.896 | 0.980 |

#### Supplemental Material

**Table S5. Estimated marginal means (EMMs) of escape response.** EMM escape-response probabilities (± SE) from fitted bias-reduced binomial GLMs for each pH–duration treatment using cluster-robust covariance matrix. Results are shown for the three analysis groups within each evaluation time point. Escape response probabilities are presented on the response scale. LCL and UCL denote the lower and upper 95% confidence limits.

| Analysis Group | Evaluation Time Point | Treatment | Probability | Std. Error | LCL | UCL |
|---|---|---|---|---|---|---|
| Adult Females (Trial 1) | 0 hpe | pH 8.0 (control) | 0.969 | 0.022 | 0.881 | 0.992 |
| | | pH10.5-1min | 0.844 | 0.134 | 0.423 | 0.975 |
| | | pH10.5-5min | 0.719 | 0.069 | 0.571 | 0.831 |
| | | pH10.5-10min | 0.719 | 0.068 | 0.571 | 0.831 |
| | 24 hpe | pH 8.0 (control) | 0.969 | 0.022 | 0.881 | 0.992 |
| | | pH10.5-1min | 0.906 | 0.061 | 0.661 | 0.98 |
| | | pH10.5-5min | 0.719 | 0.134 | 0.411 | 0.903 |
| | | pH10.5-10min | 0.906 | 0.069 | 0.661 | 0.98 |
| | 72 hpe | pH 8.0 (control) | 0.844 | 0.069 | 0.661 | 0.937 |
| | | pH10.5-1min | 0.844 | 0.134 | 0.423 | 0.975 |
| | | pH10.5-5min | 0.633 | 0.088 | 0.451 | 0.784 |
| | | pH10.5-10min | 0.833 | 0.060 | 0.682 | 0.921 |
| **Analysis Group** | **Evaluation Time Point** | **Treatment** | **Probability** | **Std. Error** | **LCL** | **UCL** |
| Juveniles (Trials 2-3**)** | 0 hpe | pH 8.0 (control) | 0.989 | 0.008 | 0.959 | 0.997 |
| | | pH10.5-1min | 0.987 | 0.009 | 0.951 | 0.997 |
| | | pH10.5-5min | 0.987 | 0.008 | 0.955 | 0.997 |
| | | pH10.5-10min | 0.961 | 0.019 | 0.900 | 0.986 |
| | 24 hpe | pH 8.0 (control) | 0.981 | 0.010 | 0.948 | 0.993 |
| | | pH10.5-1min | 0.970 | 0.019 | 0.902 | 0.991 |
| | | pH10.5-5min | 0.971 | 0.018 | 0.906 | 0.991 |
| | | pH10.5-10min | 0.849 | 0.040 | 0.754 | 0.912 |
| | 72 hpe | pH 8.0 (control) | 0.936 | 0.023 | 0.874 | 0.969 |
| | | pH10.5-1min | 0.881 | 0.033 | 0.800 | 0.932 |

#### Supplemental Material

| | | | | | | |
|---|---|---|---|---|---|---|
| | | pH10.5-5min | 0.962 | 0.019 | 0.900 | 0.986 |
| | | pH10.5-10min | 0.929 | 0.027 | 0.856 | 0.966 |
| **Analysis Group** | **Evaluation Time Point** | **Treatment** | **Probability** | **Std. Error** | **LCL** | **UCL** |
| Juveniles (Trials 4-5) | 0 hpe | pH 8.0 (control) | 0.981 | 0.011 | 0.944 | 0.994 |
| | | pH9.0-1min | 0.979 | 0.012 | 0.938 | 0.993 |
| | | pH9.0-15min | 0.980 | 0.011 | 0.942 | 0.993 |
| | | pH9.0-30min | 0.948 | 0.027 | 0.860 | 0.982 |
| | 24 hpe | pH 8.0 (control) | 0.965 | 0.013 | 0.927 | 0.984 |
| | | pH9.0-1min | 0.961 | 0.024 | 0.878 | 0.988 |
| | | pH9.0-15min | 0.980 | 0.010 | 0.948 | 0.993 |
| | | pH9.0-30min | 0.940 | 0.021 | 0.884 | 0.969 |
| | 72 hpe | pH 8.0 (control) | 0.941 | 0.021 | 0.885 | 0.971 |
| | | pH9.0-1min | 0.926 | 0.026 | 0.856 | 0.963 |
| | | pH9.0-15min | 0.964 | 0.017 | 0.909 | 0.986 |
| | | pH9.0-30min | 0.939 | 0.026 | 0.862 | 0.974 |

#### Supplemental Material

**Table S6. Wald tests for survival models.** Wald $\chi^2$ tests evaluating main effect of treatment, and trial where necessary, in the fitted bias-reduced binomial GLMs of escape response. Tests were conducted with cluster-robust covariance estimates to account for repeated measurements across evaluation time-points. DF denotes the numerator degrees of freedom. The denominator DF were infinite. Results are shown for the three analysis groups.

| **Analysis Group** | **Model Term** | **DF** | **F ratio** | **Chi square** | ***p*** |
|---|---|---|---|---|---|
| Adult Females (Trial 1) | Treatment | 3 | 0.861 | 2.583 | 0.4607 |
| **Analysis Group** | **Model Term** | **DF** | | | |
| Juveniles (Trials 2-3**)** | Treatment | 3 | 3.842 | 11.526 | 0.0092 |
| | Trial | 1 | 0.740 | 0.750 | 0.3898 |
| **Analysis Group** | **Model Term** | **DF** | | | |
| Juveniles (Trials 4-5**)** | Treatment | 3 | 0.693 | 2.079 | 0.5563 |
| | Trial | 1 | 0.025 | 0.025 | 0.8732 |

#### Supplemental Material

**Table S7. Wald tests for escape-response models.** Wald $\chi^2$ tests evaluating main and interactive effects of treatment × evaluation-time in the fitted bias-reduced binomial GLMs of escape response. Tests were conducted with cluster-robust covariance estimates to account for repeated measurements across evaluation time-points. DF denotes the numerator degrees of freedom. The denominator DF were infinite. Results are shown for the three analysis groups.

| **Analysis Group** | **Model Term** | **DF** | **F ratio** | **Chi square** | ***p*** |
|---|---|---|---|---|---|
| Adult Females (Trial 1) | Treatment | 3 | 2.980 | 8.940 | 0.0301 |
| | Time Check | 2 | 1.799 | 3.598 | 0.1655 |
| | Treatment:Time Check | 6 | 4.906 | 29.436 | 0.0001 |
| **Analysis Group** | **Model Term** | **DF** | **F ratio** | **Chi square** | ***p*** |
| Juveniles (Trials 2-3**)** | Treatment | 3 | 2.514 | 7.542 | 0.0565 |
| | Time Check | 2 | 10.969 | 21.938 | 0.0001 |
| | Trial | 1 | 0.439 | 0.439 | 0.5075 |
| | Treatment:Time Check | 6 | 2.892 | 17.352 | 0.0081 |
| **Analysis Group** | **Model Term** | **DF** | **F ratio** | **Chi square** | ***p*** |
| Juveniles (Trials 4-5**)** | Treatment | 3 | 1.581 | 4.743 | 0.1917 |
| | Time Check | 2 | 2.428 | 4.856 | 0.0882 |
| | Trial | 1 | 2.147 | 2.147 | 0.1429 |
| | Treatment:Time Check | 6 | 0.419 | 2.514 | 0.8671 |

#### Supplemental Material

**Table S8. Dunnett's tests on final survival**. Results of Dunnett's test evaluating if treatment groups varied significantly from the control, based on fitted bias-reduced binomial GLMs. Contrasts were evaluated on the logit (link) scale and exponentiated to report odds ratios. SEs and z ratios are reported on the log (odds ratio) scale. P-values are Dunnett-adjusted for multiple comparisons.

| Analysis Group | Contrast | Odds Ratio | Std. Error | Z Ratio | *p* |
|---|---|---|---|---|---|
| Adult Females (Trial 1) | (pH 10.5-1min) / pH 8.0 | 1.000 | 1.250 | 0.00 | 1.0000 |
| | (pH 10.5-5min) / pH 8.0 | 0.264 | 0.279 | -1.260 | 0.4460 |
| | (pH 10.5-10min) / pH 8.0 | 0.369 | 0.400 | -0.919 | 0.6642 |
| **Analysis Group** | **Contrast** | **Odds Ratio** | **Std. Error** | **Z Ratio** | ***p*** |
| Juveniles (Trials 2-3) | (pH 10.5-1min) / pH 8.0 | 0.508 | 0.262 | -1.313 | 0.4143 |
| | (pH 10.5-5min) / pH 8.0 | 1.310 | 0.820 | 0.432 | 0.9224 |
| | (pH 10.5-10min) / pH 8.0 | 0.288 | 0.138 | -2.604 | 0.0258 |
| **Analysis Group** | **Contrast** | **Odds Ratio** | **Std. Error** | **Z Ratio** | ***p*** |
| Juveniles (Trials 4-5) | (pH 9.0-1min) / pH 8.0 | 0.895 | 0.476 | -0.208 | 0.9828 |
| | (pH 9.0-15min) / pH 8.0 | 2.173 | 1.430 | 1.181 | 0.4953 |
| | (pH 9.0-30min) / pH 8.0 | 1.327 | 0.762 | 0.492 | 0.8987 |

#### Supplemental Material

**Table S9. Dunnett's tests on escape response**. Results of Dunnett's test evaluating if treatment groups varied significantly from the paired control group, based on fitted bias-reduced binomial GLMs. Contrasts were evaluated on the logit (link) scale and exponentiated to report odds ratios. SEs and z ratios correspond to the log (odds ratio) scale. P-values are Dunnett-adjusted for multiple comparisons. Results are shown for each analysis group within each evaluation time point.

| **Analysis Group** | **Evaluation Time** | **Contrast** | **Odds Ratio** | **Std. Error** | **Z Ratio** | ***p*** |
|---|---|---|---|---|---|---|
| Adult Females (Trial 1) | 0 hpe | (pH 10.5-1min) / pH 8.0 | 0.174 | 0.218 | -1.39 | 0.3672 |
| | | (pH 10.5-5min) / pH 8.0 | 0.082 | 0.066 | -3.11 | 0.0054 |
| | | (pH 10.5-10min) / pH 8.0 | 0.082 | 0.066 | -3.11 | 0.0054 |
| | 24 hpe | (pH 10.5-1min) / pH 8.0 | 0.312 | 0.341 | -1.064 | 0.5703 |
| | | (pH 10.5-5min) / pH 8.0 | 0.082 | 0.081 | -2.533 | 0.0314 |
| | | (pH 10.5-10min) / pH 8.0 | 0.312 | 0.341 | -1.064 | 0.5703 |
| | 72 hpe | (pH 10.5-1min) / pH 8.0 | 1.000 | 1.140 | 0 | 1 |
| | | (pH 10.5-5min) / pH 8.0 | 0.320 | 0.206 | -1.77 | 0.1909 |
| | | (pH 10.5-10min) / pH 8.0 | 0.926 | 0.627 | -0.114 | 0.995 |
| **Analysis Group** | **Evaluation Time** | **Contrast** | **Odds Ratio** | **Std. Error** | **Z Ratio** | ***p*** |
| Juveniles (Trials 2-3**)** | 0 hpe | (pH 10.5-1min) / pH 8.0 | 0.856 | 0.831 | -0.160 | 0.9900 |
| | | (pH 10.5-5min) / pH 8.0 | 0.893 | 0.852 | -0.118 | 0.9946 |
| | | (pH 10.5-10min) / pH 8.0 | 0.280 | 0.240 | -1.483 | 0.3195 |
| | 24 hpe | (pH 10.5-1min) / pH 8.0 | 0.605 | 0.506 | -0.600 | 0.8491 |
| | | (pH 10.5-5min) / pH 8.0 | 0.632 | 0.528 | -0.549 | 0.8737 |
| | | (pH 10.5-10min) / pH 8.0 | 0.107 | 0.067 | -3.561 | 0.0011 |
| | 72 hpe | (pH 10.5-1min) / pH 8.0 | 0.505 | 0.249 | -1.385 | 0.3723 |
| | | (pH 10.5-5min) / pH 8.0 | 1.719 | 1.120 | 0.830 | 0.7199 |
| | | (pH 10.5-10min) / pH 8.0 | 0.886 | 0.493 | -0.218 | 0.9811 |
| **Analysis Group** | **Evaluation Time** | **Contrast** | **Odds Ratio** | **Std. Error** | **Z Ratio** | ***p*** |
| Juveniles (Trials 4-5**)** | 0 hpe | (pH 9.0-1min) / pH 8.0 | 0.891 | 0.710 | -0.145 | 0.9918 |
| | | (pH 9.0-15min) / pH 8.0 | 0.974 | 0.784 | -0.033 | 0.9996 |

| | | | | | | |
|---|---|---|---|---|---|---|
| | | (pH 9.0-30min) / pH 8.0 | 0.354 | 0.281 | -1.310 | 0.4157 |
| | 24 hpe | (pH 9.0-1min) / pH 8.0 | 0.889 | 0.661 | -0.158 | 0.9902 |
| | | (pH 9.0-15min) / pH 8.0 | 1.783 | 1.160 | 0.885 | 0.6854 |
| | | (pH 9.0-30min) / pH 8.0 | 0.557 | 0.291 | -1.120 | 0.5340 |
| | 72 hpe | (pH 9.0-1min) / pH 8.0 | 0.784 | 0.408 | -0.467 | 0.9089 |
| | | (pH 9.0-15min) / pH 8.0 | 1.688 | 1.050 | 0.839 | 0.7141 |
| | | (pH 9.0-30min) / pH 8.0 | 0.962 | 0.562 | -0.066 | 0.9984 |

#### Supplemental Material

**Table S10. Analysis of variance (ANOVA) of routine metabolic rate (RMR).** ANOVA was used to test the main effect of elevated pH on RMR. Assay was included as a blocking factor to account for variation between two replicate respirometry assays. For the juvenile analysis group, two potential outliers were identified, and the analysis was repeated with and without these observations as a sensitivity check.

| Analysis Group | Factor | Sum Sq | DF | F value | *p* |
|---|---|---|---|---|---|
| **Adult Females** | Treatment | 0.00245 | 1 | 1.1286 | 0.3021 |
| | Assay | 0.00773 | 1 | 3.5474 | 0.0759 |
| | Residuals | 0.03923 | 18 | | |
| **Analysis Group** | **Factor** | **Sum Sq** | **DF** | **F value** | ***p*** |
| **Juveniles** | Treatment | 0.01463 | 2 | 0.4711 | 0.6285 |
| | Assay | 0.00808 | 1 | 0.5202 | 0.4760 |
| | Residuals | 0.49676 | 32 | | |
| **Analysis Group** | **Factor** | **Sum Sq** | **DF** | **F value** | ***p*** |
| **Juveniles (after outlier removal)** | Treatment | 0.01563 | 2 | 1.8834 | 0.1696 |
| | Assay | 0.00397 | 1 | 0.9577 | 0.3356 |
| | Residuals | 0.12448 | 30 | | |

### Supplemental Material

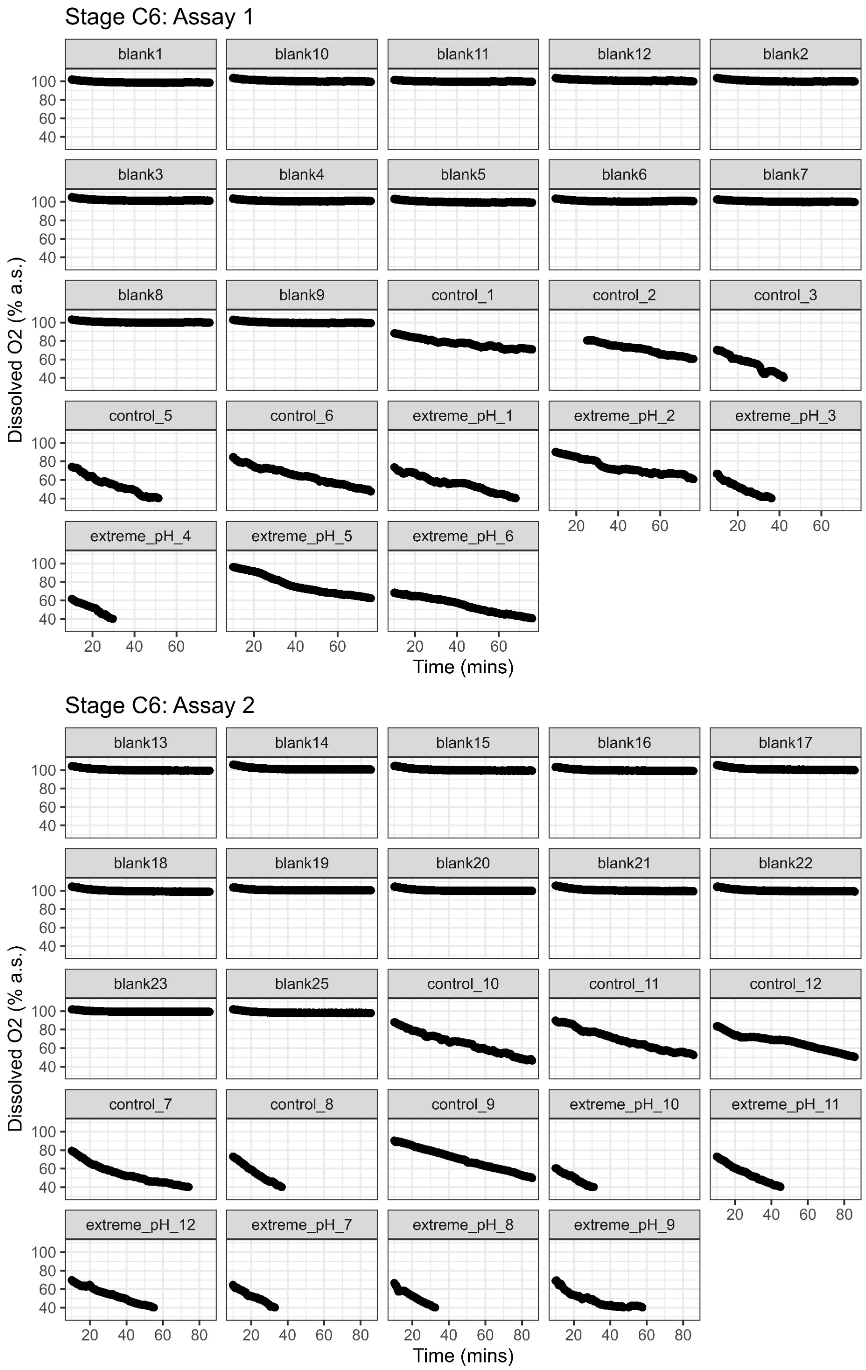


**Fig. S1.** Microplate respirometry of adult female (stage C6) *C. finmarchicus* across two replicate assays. The panels show oxygen traces (% air saturation) from individual microplate wells containing copepods and from blank wells. Wells are labeled by group (blank, control, extreme pH 10.5) and a running ID number. Each trace indicates the portion of the oxygen decline used to calculate routine metabolic rate.

### Supplemental Material

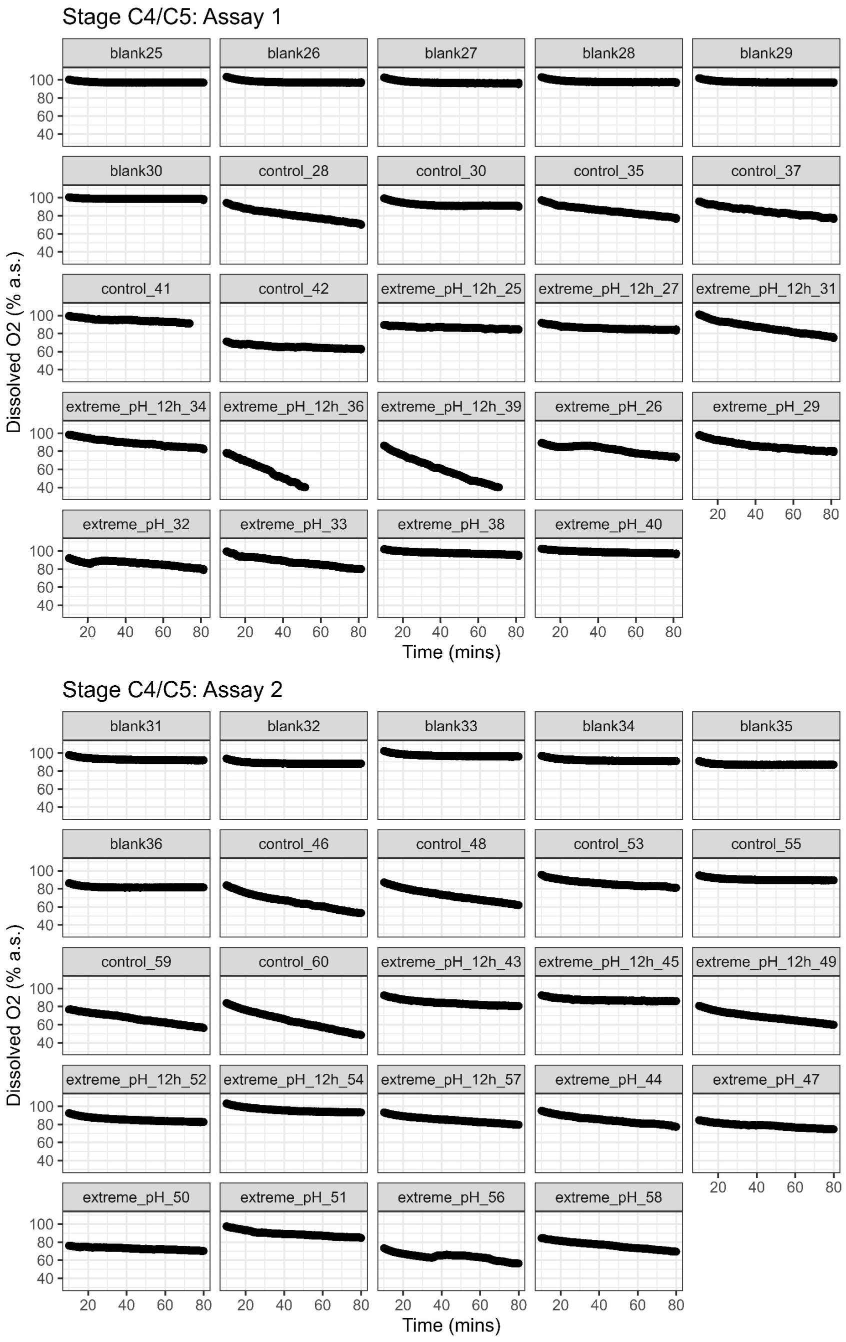


**Fig. S2.** Microplate respirometry of juvenile (stage C4-C5) *C. finmarchicus* across two replicate assays. The panels show oxygen traces (% air saturation) from individual microplate wells containing copepods and from blank wells. Wells are labeled by group (blank, control, extreme pH 10.5, extreme pH 10.5 + 12 hours post exposure) and a running ID number. Each trace indicates the portion of the oxygen decline used to calculate routine metabolic rate. Note two copepods from assay 3 were removed as outliers in the final analysis (labels: extreme_pH_12h_36, extreme_pH_12h_39).
